# Abnormal energy metabolism can alter foraging behavior in termites in different social contexts

**DOI:** 10.1101/2020.08.20.258848

**Authors:** Huan Xu, Qiuying Huang, Yongyong Gao, Jia Wu, Ali Hassan, Yutong Liu

**Affiliations:** Hubei Insect resources Utilization and Sustainable Pest Management Key Laboratory, Huazhong Agricultural University, Wuhan 430070, Hubei, China

## Abstract

Foraging behavior, as an energy-consuming behavior, is very important for collective survival in termites. How energy metabolism related to glucose decomposition and ATP production influences foraging behavior in termites is still unclear. Here, we analyzed the change in energy metabolism in the whole organism and brain after silencing the key metabolic gene isocitrate dehydrogenase (IDH) and then investigated its impact on foraging behavior in the subterranean termite *Odontotermes formosanus* in different social contexts. The *IDH* gene exhibited higher expression in the abdomen and head of *O. formosanus*. The knockdown of *IDH* resulted in metabolic disorders in the whole organism, including the impairment of the NAD^+^-IDH reaction and decreased ATP levels and glucose accumulation. The ds*IDH*-injected workers showed significantly reduced walking activity but increased foraging success. Interestingly, *IDH* downregulation altered brain energy metabolism, resulting in a decline in ATP levels and an increase in IDH activity. Additionally, the social context obviously affected brain energy metabolism and, thus, altered foraging behavior in *O. formosanus*. We found that the presence of predator ants increased the negative influence on the foraging behavior of ds*IDH*-injected workers, including a decrease in foraging success. However, an increase in the number of nestmate soldiers could provide social buffering to relieve the adverse effect of predator ants on worker foraging behavior. Our orthogonal experiments further verified that the role of the *IDH* gene as an inherent factor was dominant in manipulating termite foraging behavior compared with external social contexts, suggesting that energy metabolism, especially brain energy metabolism, plays a crucial role in regulating termite foraging behavior.

**Author summary:** Foraging behavior plays a key role in collective survival in social insects, as found in termites. Worker termites are responsible for foraging duty and exhibit large foraging areas and long foraging distances, so they need to consume much energy during foraging. It is well established that energy can influence insect behaviors. However, how energy metabolism affects foraging behavior in termites remains unknown. Here, we found that the downregulation of the conserved metabolic gene *IDH* impaired whole-organism and the brain energy metabolism and further altered foraging behavior, resulting in decreased walking activity but increased foraging success in the termite *O. formosanus*, which is an important insect pest damaging embankments and trees in China. Additionally, the social context affected brain energy metabolism and obviously changed foraging behavior in *O. formosanus*, causing a decline in foraging success in the absence of nestmate soldiers and the presence of predator ants. However, the increasing number of nestmate soldiers strengthened social buffering to relieve the negative effect of predator ants on worker foraging behavior. Our findings provide new insights into the underlying molecular mechanism involved in modulating the sophisticated foraging strategy of termites in different social contexts from the perspective of energy metabolism.

## Introduction

All behaviors consume energy (to various degrees), and some also facilitate energy acquisition, such as foraging behavior, which presents a cost-benefit trade-off between growth and survival in organisms [1, 2]. The foraging behavior of social insects is an extremely complex process involving the self-organization of a large number of individuals to collect foods from various sources [3, 4]. For example, the honey bee *Apis mellifera* recruits additional foragers to valuable resources by dancing to signal the location of the resources to their nestmates [5]. In seed-harvesting ants of *Pogonomyrmex* spp., a mixture of individual and group foraging is the most common foraging strategy [6]. A group of foraging workers utilizes clay to build structural supports to increase foraging resources in the termite *Coptotermes acinaciformis* [7]. Therefore, social insects prefer to employ highly effective and energy-saving foraging strategies to obtain sufficient food and thereby fulfil the energy requirements of their colonies.

A large body of research is focused on the relationship between energy metabolism and behavior in both vertebrates and invertebrates [1, 8–10], revealing that the brain is a major regulator of behavior and metabolic processes [11, 12]. For instance, in adult *Drosophila*, the knockdown of glycolytic enzymes specifically in the glial cells of the nervous system in the brain leads to severe locomotor deficits [13]. The regulation of behavior is affected by the nerves and requires large amounts of energy in the brains of the cricket *Gryllus campestris* L. and the ant *Pheidole rhea* [14, 15]. It seems that the basic energy metabolism related to neuromodulation can explain the relationship between energy metabolism and behavior [16, 17]. However, the role of energy metabolism in regulating social behaviors is still unclear.

Complex social contexts can affect foraging behavior in social insects. Considerable research has shown that predation risk seriously affects animal activity and foraging behavior [18–22]. Biological individuals may make collective decisions in intricate social contexts [23] and then respond to the presence of predators by showing a range of behavioral and physiological changes to reduce the likelihood of injury or death [24]. For instance, cryptic termites can change their foraging strategy by eavesdropping on vibrational cues from the footsteps of predatory ants to limit predation risk [25, 26]. Honey bees avoid flowers with crab spiders and flowers that have recently held spiders during foraging [27]. Therefore, predation pressure from predators is an important factor impacting foraging behavior in social insects. In addition, social information provided by nestmates can also change individual foraging decisions [28]. In social insects, soldiers play a vital role in coping with the risk of external predation by sending out warning signals in a special way so that the other nestmates can avoid risks in a limited time through nestmate recognition during the process of foraging [29–31]. Social buffering is involved in the ability of neighboring individuals in a colony to reduce the negative impact of stressors on other individuals in a wide range of vertebrates [32]. For example, in the termite *Reticulitermes flavipes*, the presence of nestmate soldiers can change the ability of workers to cope with the competition risk imposed by conspecific non-nestmates [24]. However, the changes in behavioral characteristics are unclear when nestmate soldiers provide social buffering to foraging termites with different energy levels in the presence of predator ants.

Theoretically, both inherent genes and external social contexts can influence foraging behavior in insects. The *foraging* gene (*for*), encoding a cGMP-dependent protein kinase (PKG), is related to the foraging behavior of several social and solitary insects, including the lower termite *R. flavipes* [33], the fruit fly *Drosophila melanogaster* [34, 35], the honey bee *Apis mellifera* [36], the bumblebee *Bombus ignites* [37], the red harvester ant *Pogonomyrmex barbatus* [38], the ant *Pheidole pallidula* [39] and the fire ant *Solenopsis invicta* [40]. In solitary *D. melanogaster*, *foraging* genes can influence a number of behavioral traits, which can also be modified by the social context [41–43]. However, how the complex interactions of inherent genes and social contexts give rise to variations in foraging behavior in termites is still unclear.

The subterranean termite *Odontotermes formosanus* (Shiraki), which is a fungus-cultivating higher termite, can construct large underground cavities and damage many kinds of trees [44, 45]. Workers exhibit large foraging areas and long foraging distances in the termite *O. formosanus* [46]. They need to build mud lines during their foraging to avoid adverse environmental factors and increase foraging resources [47–49]. Thus, sufficient energy reserves appear to be an important factor affecting the likelihood of foraging success in termites [50]. Isocitrate dehydrogenase (IDH) is an important rate-limiting enzyme that catalyzes the decarboxylation of isocitrate to alpha-ketoglutarate in the tricarboxylic acid cycle (TCA) and generates nicotinamide-adenine dinucleotide (NADH), which feeds into oxidative phosphorylation to generate adenosine triphosphate (ATP) [51]. A recent study showed that *IDH* downregulation can disrupt active immunization against the entomopathogenic fungus *Metarhizium anisopliae* in the termite *R. chinensis* [52]. In this study, we wanted to determine whether *IDH* gene knockdown impairs energy metabolism involved in glucose decomposition and ATP production and further affects foraging behavior in termites. Here, we cloned the *IDH* gene encoding the α subunit of IDH and observed its expression pattern in *O. formosanus*. Subsequently, we silenced the *IDH* gene and explored the effect of energy metabolism alterations in the whole organism and brain on the foraging behavior of *O. formosanus*. Moreover, we carried out orthogonal experiments to analyze the changes in *IDH*-mediated foraging behavior in different social contexts (with or without *Leptogenys kitteli* predator ants or/and nestmate soldiers) according to four experimental parameters (velocity, distance moved, frequency and cumulative duration in food zones) in *O. formosanus*.

## Results

### Cloning, expression and RNAi knockdown efficiency of the *IDH* gene in *O. formosanus*

The complete open reading frame (ORF) of *IDH* was 1,074 bp and encoded a predicted protein of 357 amino acids (aa), which is highly conserved in social organisms (Fig 1A). The prediction of the secondary structure of the IDH protein demonstrated that the amino acid sequence contained 47.90% alpha-helices, 10.36% extended strands, 6.16% beta-turns, and 35.57% random coils (Fig 1B). The analysis of the conserved domains with the NCBI tool indicated that IDH presented the specific hit Iso_dh (26-350 aa) (Fig 1B). The evolutionary tree (Fig 1C) showed that *IDH* from *O. formosanus* was clustered with that from the German cockroach, *Blattella germanica*, and two termite species, *Zootermopsis nevadensis* and *Cryptotermes secundus*.

**Fig 1.**
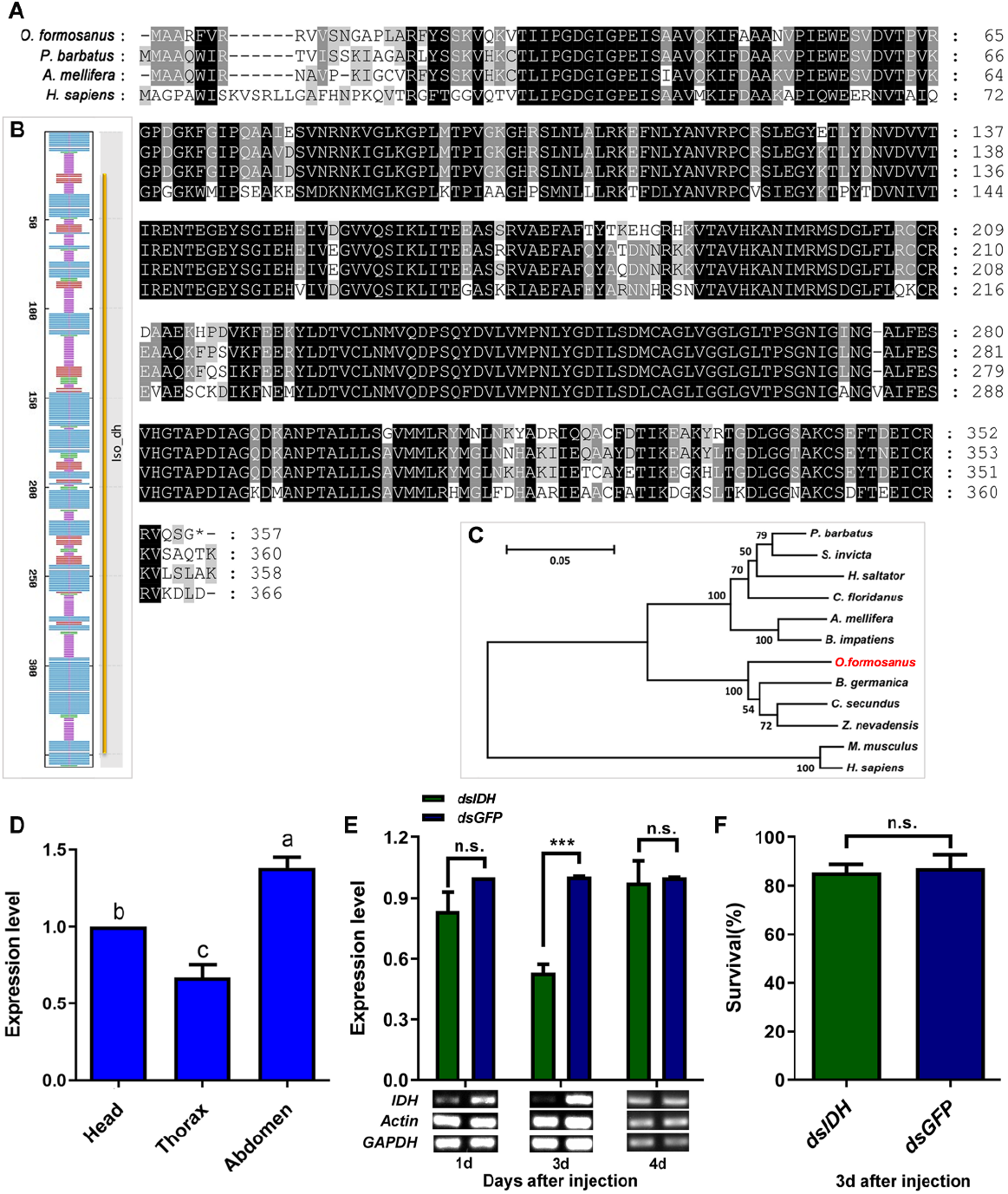
Cloning, expression and RNAi efficiency of the *IDH* gene in *O. formosanus*. (A) Multiple alignments of the amino acid sequences deduced for IDH in *O. formosanus* and three other social species. Identical residues were shaded in black. Dashed lines indicated the gaps. (B) Secondary structure prediction of the amino acid sequence of IDH and conserved domains of *IDH*. Random coils were shown in purple, alpha-helixes were shown in blue, beta-turns were shown in green, and extended strands were shown in red. The gene showed an Iso_dh specific hit (26-350). (C) Phylogenetic relationships of the IDH proteins from *O. formosanus* and 11 other social species. (D) Expression patterns of *IDH* in the head, thorax and abdomen of workers. (E) Expression level and semiquantitative detection of *IDH* 1, 3 and 4 days after dsRNA injection in workers; (F) Survival 3 d after dsRNA injection in workers. Different lowercase letters in a column indicate significant differences by Tukey’s HSD test. ***, *p* < 0.001, n.s. means no significant difference.

Our results showed that *IDH* expression in the abdomen and head was significantly higher than that in the thorax (Fig 1D, Tukey’s HSD test, F = 30.380, df = 2,15, *p* < 0.001, n = 6). The expression level of *IDH* was significantly decreased by 47.44% 3 d after the introduction of RNAi (Fig 1E, paired *t*-test, *t* = −11.055, df = 8, *p* < 0.001, n = 9). Semiquantitative RT-PCR analysis also showed that the expression of *IDH* was successfully inhibited (Fig 1E). In addition, there were no significant differences in survival between ds*IDH*-injected individuals and ds*GFP*-injected individuals (Fig 1F, paired *t*-test, *t* = −0.419, df = 5, *p* = 0.692, n = 6).

### *IDH* knockdown reduced the energy supply but increased glucose accumulation

The IDH enzyme is an important rate-limiting enzyme in the TCA and can catalyze the decarboxylation of isocitrate to alpha-ketoglutarate [51]. Therefore, we silenced *IDH* to investigate the changes in the NAD^+^-IDH reaction in the TCA and the level of glucose 3 d after the injection of ds*IDH*. The results showed that IDH activity (Fig 2A, Wilcoxon test, Z = −2.023, *p* = 0.043, n = 6), ATP levels (Fig 2B, Wilcoxon test, Z = −2.666, *p* = 0.008, n = 9) and NADH levels (Fig 2C, Wilcoxon test, Z = −2.201, *p* = 0.028, n = 6) were significantly decreased in ds*IDH*-injected termites compared to ds*GFP*-injected termites. However, the glucose level was significantly increased compared with that in ds*GFP*-injected termites (Fig 2D, Wilcoxon test, Z = −2.192, *p* = 0.028, n = 9).

**Fig 2.**
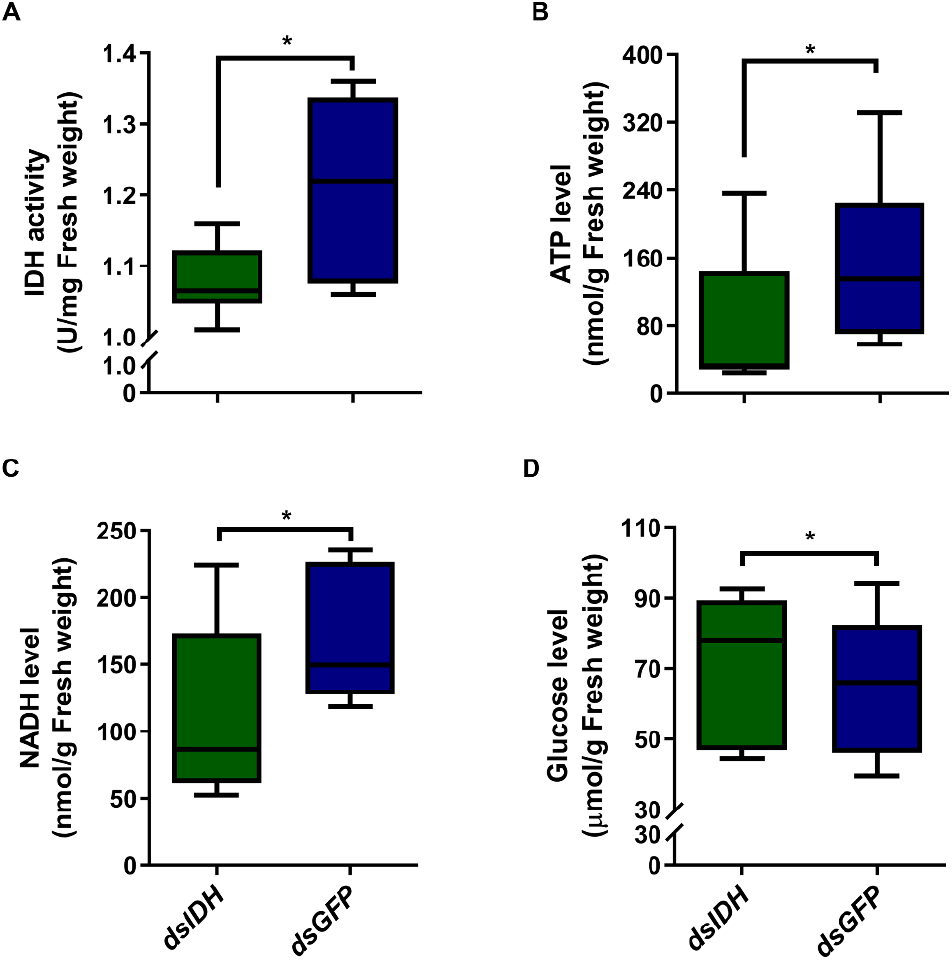
Alterations in energy metabolism after silencing *IDH* in *O. formosanus*. (A) IDH activity in workers 3 d after ds*IDH* injection; (B) ATP levels in workers 3 d after ds*IDH* injection; (C) NADH levels in workers 3 d after ds*IDH* injection; (D) Glucose levels in workers 3 d after ds*IDH* injection. \**p* < 0.05.

### Silencing *IDH* decreased walking activity but increased foraging success

To investigate the relationship between energy metabolism and foraging behavior, we silenced *IDH* by injecting ds*IDH* and then observed the phenotypic changes in the foraging behavior (Fig 3A) of *O. formosanus*. We found that the foraging trajectories of ds*GFP*-injected individuals were denser than those of ds*IDH*-injected individuals (Fig 3B). Both the distance moved (Fig 3C, paired *t*-test, *t* = −2.502, df = 9, *p* = 0.034, n = 10) and velocity (Fig 3D, paired *t*-test, *t* = −2.448, df = 9, *p* = 0.037, n = 10) were significantly decreased in ds*IDH*-injected individuals compared to ds*GFP*-injected individuals. However, the frequency (Fig 3E, paired *t*-test, *t* = 2.890, df = 9, *p* = 0.018, n = 10) and the cumulative duration (Fig 3F, paired *t*-test, *t* = 2.344, df = 9, *p* = 0.044, n = 10) in food zones were significantly increased in ds*IDH*-injected individuals compared with ds*GFP*-injected individuals.

**Fig 3.**
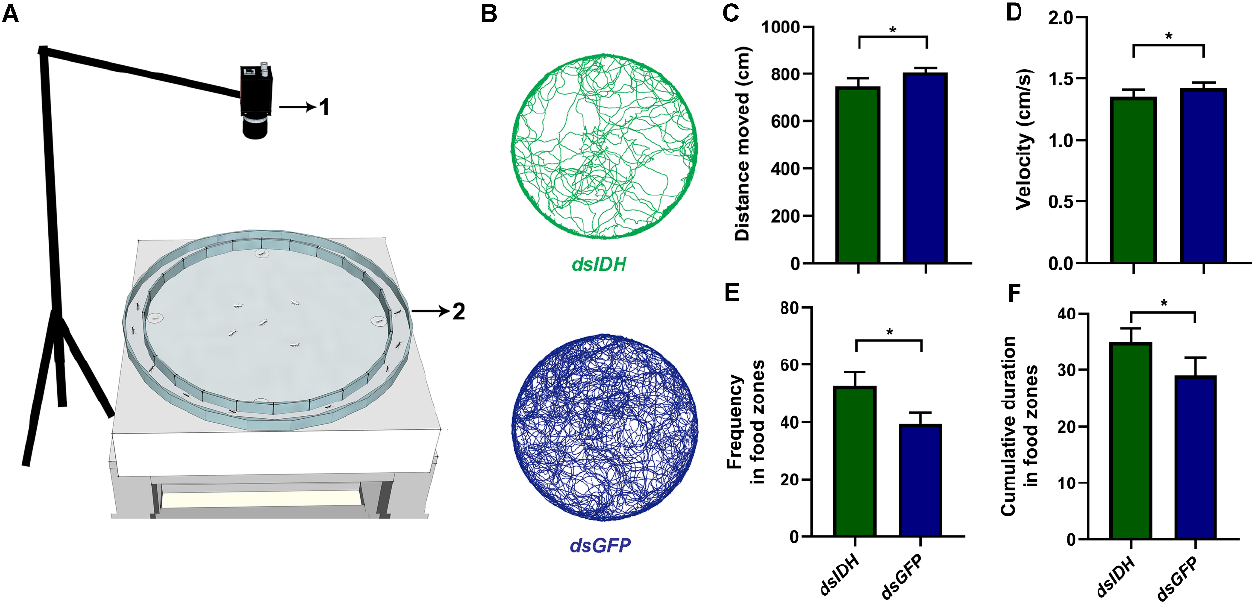
Silencing *IDH* decreased walking activity but increased foraging success in *O. formosanus*. (A) Behavioral apparatus (1, camera; 2, testing arena). (B) Foraging trajectories of ds*IDH*-injected (deep green color) and ds*GFP*-injected (deep blue color) workers. (C) The distance moved of ds*IDH*-injected and ds*GFP*-injected workers. (D) The velocity of ds*IDH*-injected and ds*GFP*-injected workers. (E) The frequency in food zones of ds*IDH*-injected and ds*GFP*-injected workers. (F) The cumulative duration in food zones of ds*IDH*-injected and ds*GFP*-injected workers. * *p* < 0.05.

### The effect of *IDH* on brain energy metabolism in different social contexts

We successfully obtained the brain tissue of worker termites (Fig 4A) and found that there were a large number of tracheae in the brain of *O. formosanus*. Immunocytochemistry analysis of synapsin revealed the major medullary structures in the workers’ brains (Fig 4B). Our results showed that *IDH* expression in the worker brain was significantly decreased 3 d after ds*IDH* injection (Fig 4C, Wilcoxon test, Z = −2.201, *p* = 0.028, n = 6). In three different social contexts, IDH activity in the brains of ds*IDH*-injected worker individuals was significantly higher than that in ds*GFP*-injected worker individuals, respectively (Fig 4D, Wilcoxon test, Z_1_ = −1.992, *p_1_* = 0.046, n_1_ = 6; Z_2_ = −2.201, *p_2_* = 0.028, n_2_ = 6; Z_3_ = −2.201, *p_3_* = 0.028, n_3_ = 6), while ATP levels in the brains of ds*IDH*-injected worker individuals were significantly lower than that in ds*GFP*-injected worker individuals, respectively (Fig 4E, Wilcoxon test, Z1 = −2.201, *p_1_* = 0.028, n_1_ = 6; Z_2_ = −2.201, *p_2_* = 0.028, n_2_ = 6; Z_3_ = −2.201, *p_3_* = 0.028, n_3_ = 6). Additionally, we found that IDH activity in the worker brain was significantly higher in the social context with ants and soldiers than that in the social context without ants and soldiers (Fig 4D, Wilcoxon test, ds*IDH*: Z = −1.992, *p* = 0.046, n = 6; ds*GFP*: Z = −2.201, *p* = 0.028, n = 6). However, ATP levels in the worker brain were significantly lower in the social context with ants and soldiers than that in the social context without ants and soldiers (Fig 4E, Wilcoxon test, ds*IDH*: Z = −2.201, *p* = 0.028, n = 6; ds*GFP*: Z = −2.201, *p* = 0.028, n = 6).

**Fig 4.**
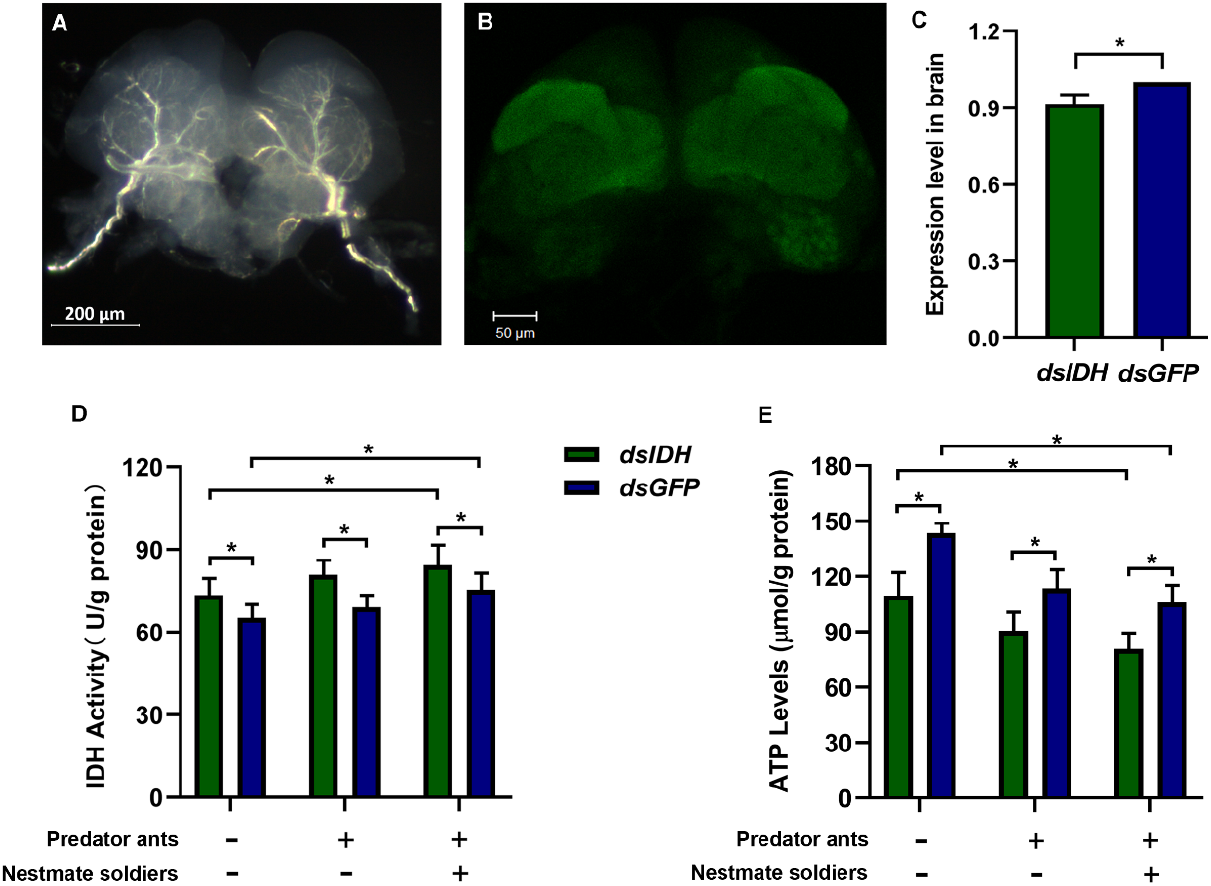
Effect of *IDH* silencing on brain energy metabolism in *O. formosanus* in different social contexts. (A) Full brain tissue of workers; (B) Confocal image of the worker brain; (C) Expression level of *IDH* in the brains of workers 3 d after ds*IDH* injection; (D) IDH activity in the brains of ds*IDH*-injected and ds*GFP*-injected workers in different social contexts. (E) ATP levels in the brains of ds*IDH*-injected and ds*GFP*-injected workers in different social contexts. * *p* < 0.05.

### The influence of *IDH* on foraging behavior in different social contexts

To further explore the impact of *IDH* on the foraging behavior of *O. formosanus* in different social contexts, we set up nine treatment groups with or without predator ants or/and nestmate soldiers using orthogonal experimental design L_9_ (3^3^) (Table 1, Fig 5A-D). Our results showed that the foraging trajectories of *O. formosanus* in the nine treatment groups varied, which was associated with the status of worker energy metabolism and different social contexts (Fig 5E). The foraging trajectories of water-injected workers became increasingly complex and chaotic along with the increase in the number of ants (from Treatment 1 to 3). The foraging trajectories of ds*GFP*-injected workers were similar to those of water-injected workers (from Treatment 4 to 6). However, the foraging trajectories of ds*IDH*-injected workers exhibited less complexity and lower flexibility when the number of ants increased (from Treatment 7 to 9).

**Table 1.** The features of foraging behavior in *O. formosanus* under different RNAi treatments and social contexts

| Treatment | Factors (levels) |  |  | Results |  |  |  |
| --- | --- | --- | --- | --- | --- | --- | --- |
| number | Gene | Ant<br>number | Soldier<br>number | Velocity<br>(cm/s) | Distance moved<br>(cm) | Frequency<br>in food zones | Cumulative duration<br>in food zones (s) |
| 1 | *water* | 0 | 0 | 1.62±0.10a | 900.31±56.41a | 46.60±2.96a | 41.86±1.85a |
| 2 | *water* | 10 | 1 | 1.60±0.12a | 879.71±74.34ab | 41.53±4.02ab | 38.92±2.18ab |
| 3 | *water* | 20 | 2 | 1.59±0.11a | 856.93±61.35ab | 39.52±3.64abc | 36.61±2.27ab |
| 4 | *dsGFP* | 0 | 1 | 1.45±0.13ab | 799.60±63.44abc | 39.63±4.86abc | 38.16±2.21ab |
| 5 | *dsGFP* | 10 | 2 | 1.33±0.09ab | 707.54±53.08abc | 27.35±4.14bc | 29.72±4.11ab |
| 6 | *dsGFP* | 20 | 0 | 1.57±0.12a | 859.54±70.34ab | 38.70±4.47abc | 36.03±3.22ab |
| 7 | *dsIDH* | 0 | 2 | 1.09±0.09b | 583.60±48.83c | 30.60±3.97abc | 34.30±3.10ab |
| 8 | *dsIDH* | 10 | 0 | 1.22±0.07ab | 630.35±48.70bc | 22.67±2.75c | 27.72±2.46b |
| 9 | *dsIDH* | 20 | 1 | 1.31±0.10ab | 663.02±59.63abc | 31.22±3.29abc | 29.13±2.88b |
| Velocity (cm/s) |  |  |  |  |  |  |  |
| *R* | 0.40 | 0.11 | 0.13 |  |  |  |  |
| *F* | 11.408*** | 1.018 | 1.582 |  |  |  |  |
| Distance moved (cm) |  |  |  |  |  |  |  |
| *R* | 253.33 | 53.96 | 80.71 |  |  |  |  |
| *F* | 13.925*** | 0.622 | 1.543 |  |  |  |  |
| Frequency in food zones |  |  |  |  |  |  |  |
| *R* | 14.39 | 8.43 | 4.97 |  |  |  |  |
| *F* | 10.619*** | 3.850* | 1.339 |  |  |  |  |
| Cumulative duration in food zones |  |  |  |  |  |  |  |
| *R* | 8.75 | 4.56 | 1.86 |  |  |  |  |
| *F* | 7.455*** | 3.671* | 0.404 |  |  |  |  |

*R* indicates the impact degree of the three factors on the features of foraging behavior obtained from the $L_9 (3^3)$ orthogonal experiment. A higher *R* value for a factor suggests a stronger effect on foraging behavior. The data in the table are the mean ± SEM, and different letters in a column indicate significant differences by Tukey's HSD test ( $p < 0.05$ , $n = 12$ ). Asterisks indicate significant differences by GLM (\*\*\* $< 0.001$ , $0.01 < * < 0.05$ ).

**Fig 5.**
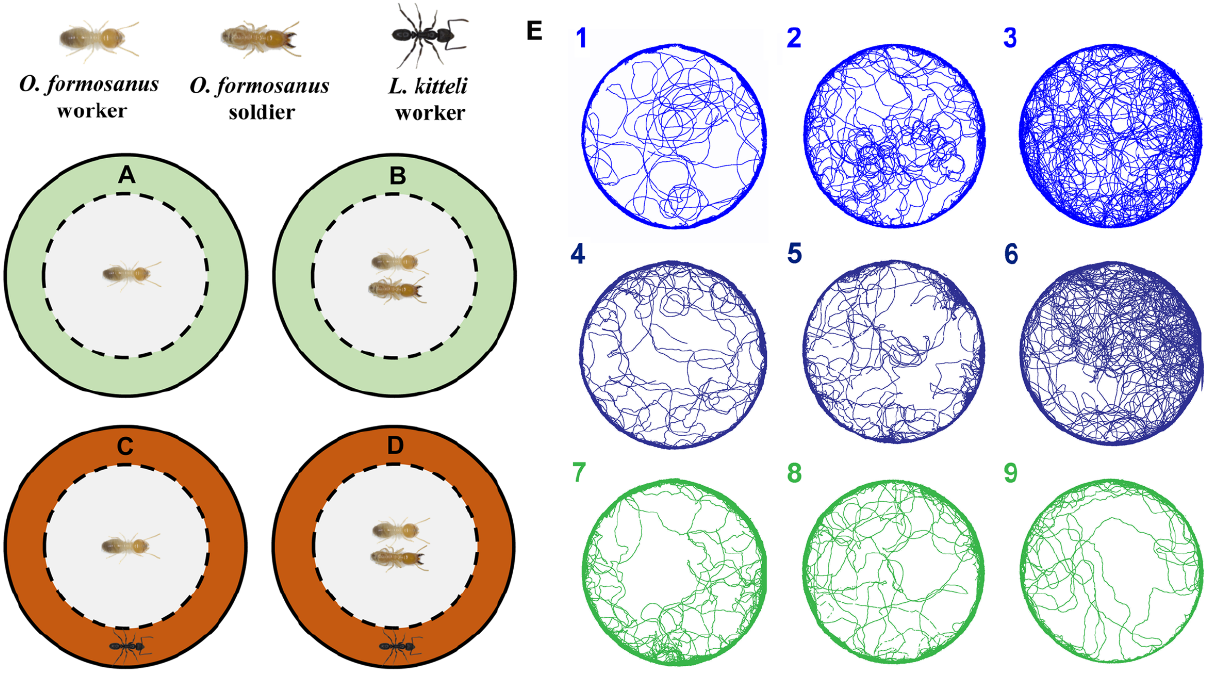
Foraging trajectories after *IDH* silencing in *O. formosanus* in different social contexts. Workers without (A) or with (B) soldiers were confined in the inner ring (in gray) without ants in the outer ring (areas in light green). The wall of the center dish was cut vertically to make 1 mm-wide silts. Predation risk was perceived by the antennation of workers of the ant *L. kitteli* through the slits. The predator ants were placed in the outer ring (areas in deep orange) without (C) or with (D) soldiers. (E) Representative maps of foraging trajectories in the water-injected (in blue), ds*GFP*-injected (in deep blue) and ds*IDH*-injected (in deep green) workers. Numbers in the upper left corner of each trajectory map indicate different treatment numbers in the orthogonal experiments L_9_ (3^3^).

There were significant differences in the phenotypic features of foraging behavior among the nine treatment groups in worker termites (Table 1). The velocity in Treatment 7 was much lower than that in Treatments 1, 2, 3 and 6 (Tukey’s HSD test, F = 3.433, df = 8, 99, *p* = 0.002). At the same time, the distance moved in Treatment 7 was extremely significantly shorter than that in Treatment 1 and significantly shorter than that in Treatments 2, 3 and 6 (Tukey’s HSD test, F = 3.964, df = 8, 99, *p* < 0.001). In addition, the distance moved in Treatment 8 was significantly shorter than that in Treatment 1 (Tukey’s HSD test, F = 3.964, df = 8, 99, *p* < 0.001). However, the frequency in food zones in Treatment 5 was significantly less than that in Treatment 1, and the frequency in food zones in Treatment 8 was extremely significantly lower than that in Treatment 1 and was markedly less than that in Treatment 2 (Tukey’s HSD test, F = 4.027, df = 8, 99, *p* < 0.001). Additionally, the cumulative duration of visits in food zones in Treatment 8 was significantly shorter than that in Treatment 1, and the cumulative duration of visits in food zones in Treatment 9 was also extremely significantly shorter than that in Treatment 1 (Tukey’s HSD test, F = 3.095, df = 8, 99, *p* = 0.004).

The impact degrees of three factors on the velocity and distance moved were as follows: *IDH* expression > soldier number > ant number, but the impact degrees of the three factors on the frequency and cumulative duration in food zones were as follows: gene expression > ant number > soldier number. Among these factors, *IDH* expression showed a significant influence on the velocity and distance moved, and frequency and cumulative duration in food zones (GLM, all *p* < 0.001). Additionally, ant number exerted a significant influence on the frequency and cumulative duration in food zones (GLM, *p_1_* = 0.024; *p_2_* = 0.029).

## Discussion

The abnormal energy metabolism of the whole organism mediated by the IDH gene seriously altered the foraging behavior of *O. formosanus*. The *IDH* gene, associated with energy metabolism, was highly expressed in the abdomen and head of foraging workers, which supports the notion that the movement requires much energy [53], particularly for foraging behavior, which requires integration across different sensory modalities in worker termites. In our study, silencing *IDH* disrupted mitochondrial metabolism, including the disruption of the NAD^+^-IDH reaction in the TCA cycle and the reduction of ATP levels, NADH levels and IDH activity (Fig 2A, B and C), which caused decreased walking activity, including shorter collective distances moved and slower velocities. Additionally, the trajectories of worker termites showed lower mobility (Fig 3B). These results suggest that mitochondrial metabolism plays a vital role in the regulation of walking activity. Normally, glucose can be decomposed and then produce ATP through the glycolysis pathway and TCA cycle [51]. However, a significant increase in glucose content was found in this study after silencing *IDH*, which was also recently described in association with the downregulation of human *IDH3α* [54] and knockdown of termite *IDH* [52]. Our results indicated that silencing *IDH* disrupted the process of glucose metabolism and resulted in the accumulation of glucose [52]. Surprisingly, knocking down *IDH* increased the frequency and cumulative duration in food zones within a certain amount of foraging time, suggesting that termite colonies could ensure the food supply by increasing foraging success [35] and prolonging the time of food acquisition to reduce the negative effects of energy deficiency. The changes in the energy metabolism of the whole organism caused by *IDH* knockdown in this study provided direct evidence of the importance of the rate-limiting enzyme IDH gene in regulating metabolic homeostasis and energy supply, which is closely related to foraging behavior in termites.

*IDH* expression levels might change the energy metabolism pathway of the termite brain, which is energetically demanding as well as a key regulator of energy metabolism [55]. In this study, *IDH* knockdown significantly reduced ATP levels in the brains of worker termites but dramatically improved IDH activity. These results suggested that the production efficiency of ATP decreased. Usually, glucose is metabolized via glycolysis and the TCA cycle to generate sufficient ATP through the process of oxidative phosphorylation [56]. However, *IDH* downregulation disrupts the TCA cycle and thereby reduces the generation of ATP. On the other hand, it has been demonstrated that a decrease in the expression of *IDH3α* converts the metabolic mode from oxidative phosphorylation to aerobic glycolysis in cancer-associated fibroblasts [54]. However, glycolysis is far less efficient than the TCA cycle coupled to oxidative phosphorylation in producing ATP [56]. Therefore, the decreasing ATP levels in the brain of ds*IDH*-injected termites suggests that the brain might choose an inefficient way of producing ATP, such as aerobic glycolysis, resulting in a low efficiency of ATP generation. Actually, ATP is necessary to maintain homeostasis and cell survival, and the loss of intracellular ATP may result in cell necrosis or apoptosis [56, 57]. In addition, energetic constraints play a major role in neural plasticity and brain health [58]. Thus, an increase in IDH activity might reduce the damage caused by ATP deficiency in ds*IDH*-injected termites.

Energy metabolism in the termite brain is not only impacted by *IDH* expression levels but is also influenced by the social context. Some researchers have suggested that social information can change gene expression in the brain to influence behaviors in social insects [59]. We also found that social information could affect foraging behavior by changing brain energy metabolism in termites. When worker termites are exposed to predation risks during foraging, they may exhibit an aggressive state and consume much energy [29]. At this moment, ATP levels decreased remarkedly, but the IDH activity increased significantly in the brains of ds*IDH*- and ds*GFP*-injected foragers in the social context with predators, suggesting that the degree of aerobic glycolysis might be enhanced, as reported for the metabolic pathway in the brain of aggressive honey bees [57]. In other words, the brains of worker termites may prefer an inefficient but faster pathway for ATP production [56] to meet the high energy demands of foraging behavior when they are threatened by predators. On the other hand, the addition of nestmate soldiers during foraging may enhance the group cognition involved in the emergent aggregation behavior of workers and social networks [60, 61], which is energetically costly in brains [12]. Therefore, the decrease in ATP became more significant, and the increase in IDH activity was greater when soldiers and ants were present concurrently. In addition, the social context may regulate the expression of genes associated with encoding neuropeptides in the brain, such as gonadotropin-releasing hormone (GnRH), vasotocin (VT) and vasopressin (VP) [62, 63]. The sensory and higher-order integrative processing mechanisms that are used when insects behave in complex social contexts are also associated with nervous systems of the brain [64], and the energetic basis of these behaviors is a bridge between behavioral ecology and neuroscience [16]. Moreover, intermediates of energy metabolism can impact the concentration of neurotransmitters by regulating the synthesis of precursors such as alpha-ketoglutarate, which is the precursor of the excitatory neurotransmitter glutamate produced via the TCA cycle [57, 65]. Thus, the changes in energy metabolism in the brain caused by *IDH* downregulation and social context might alter the energetic and nervous system bases of foraging behavior in termites.

The abnormal energy metabolism in the whole organism and the brain caused by *IDH* downregulation and the social context could modulate foraging behavior in termites. Silencing *IDH* reduced the walking activity of worker termites and simplified their foraging trajectory (Treatments 7, 8 and 9 in Table 1). This reduction in walking activity decreases significantly when the number of nestmate soldiers was increased but no ants were present (Treatment 7 in Table 1) compared to the water- or ds*GFP*-injected workers, with normal energy metabolism (Treatments 1, 2, 3 and 6 in Table 1), indicating that worker termites may choose a more profitable foraging strategy, such as reducing their foraging area, to relieve the energy shortage [66, 67] because more energy may be needed to cover a larger movement space [68, 69]. The foraging success of ds*IDH*-injected workers significantly increased (Fig 3E and F), while it decreased sharply when predator ants were present without nestmate soldiers (Treatments 8 in Table 1), indicating that predator ants seriously affect foraging behavior and that the absence of nestmate soldiers means that foraging workers face a greater risk of predation if worker termites forage for a long time and feed on a large amount of food when the energy supply is lacking [2]. Although energy deficiency caused the foraging workers to significantly reduce the cumulative duration of visits in food zones, the increasing number of soldiers enhanced the frequency in food zones appropriately (Treatment 9 in Table 1), illustrating that soldiers can provide social buffering to cover the damage caused by the increase in ants during foraging [24, 32, 70].

However, the foraging trajectory of worker termites with normal energy metabolism became complex when ants were present (Treatments 3 and 6 in Table 1), indicating that the increase in ants led to more purposeless walking, which is energetically demanding [19, 22]. In addition, the walking activity and foraging success of the worker termites showed no significant changes as the number of soldiers was increased together with the number of ants (Treatments 1, 2 and 3 in Table 1). Therefore, the presence of nestmate soldiers is crucial to reduce the purposeless walking of foraging workers and relieve the predation pressure from ants to help foragers to accurately find food sites [24]. Even though the increase in nestmate soldiers could decrease the purposeless walking caused by a small number of ants, foraging success still decreased when the number of soldiers increased (Treatment 5 in Table 1). In addition, orthogonal experiments showed that *IDH* expression extremely significantly impacted both walking activity and foraging success in foraging workers (Table 1), but predator ants only significantly influenced foraging success in foraging workers (Table 1), strongly suggesting that the inherent gene plays the dominant role in modulating foraging behavior in termites compared to the external social context [35].

In summary, we confirmed that the metabolic gene *IDH* was an important regulator of foraging behavior in termites, which can be influenced by different social contexts. *IDH* downregulation reduced ATP levels and IDH activity, and disrupted the NAD^+^-IDH reaction in the TCA in the whole organism but increased IDH activity and decreased ATP levels in the brains of termites (Fig 6). These alterations of the energetic metabolism resulted in decreased walking activity and increased foraging success, suggesting the important role of energy metabolism in the regulation of foraging behavior (Fig 6). In addition, social contexts can also change termite foraging behavior by adding predation risk from predator ants and providing social buffering from nestmate soldiers. The foraging success of ds*IDH*-injected workers decreased significantly with the addition of predator ants. The number of nestmate soldiers could improve the foraging success of ds*IDH*-injected workers by providing social buffering to relieve the negative effect of predator ants. Regardless of how the external social context changed, the impact level of the inherent IDH gene was always highest on termite foraging behavior (Table 1), which further demonstrates that the inherent gene plays a dominant role in regulating foraging behavior in termites. Hence, the present study strongly supports the hypothesis that abnormal energy metabolism can alter foraging behavior in termites in different social contexts. Certainly, whether the change in brain energy affects neural functions that may be related to foraging behavior needs to be resolved in the future. This study enriches the understanding of the energetic basis and strategy of foraging behavior in termites by verifying an energetic alteration impacting foraging behavior from the whole-organism level to the brain level in different social contexts, which will be beneficial for clarifying the role of the energy supply, allocation and consumption in determining the diversity, plasticity and evolution of social behaviors in insect societies.

**Fig 6.**
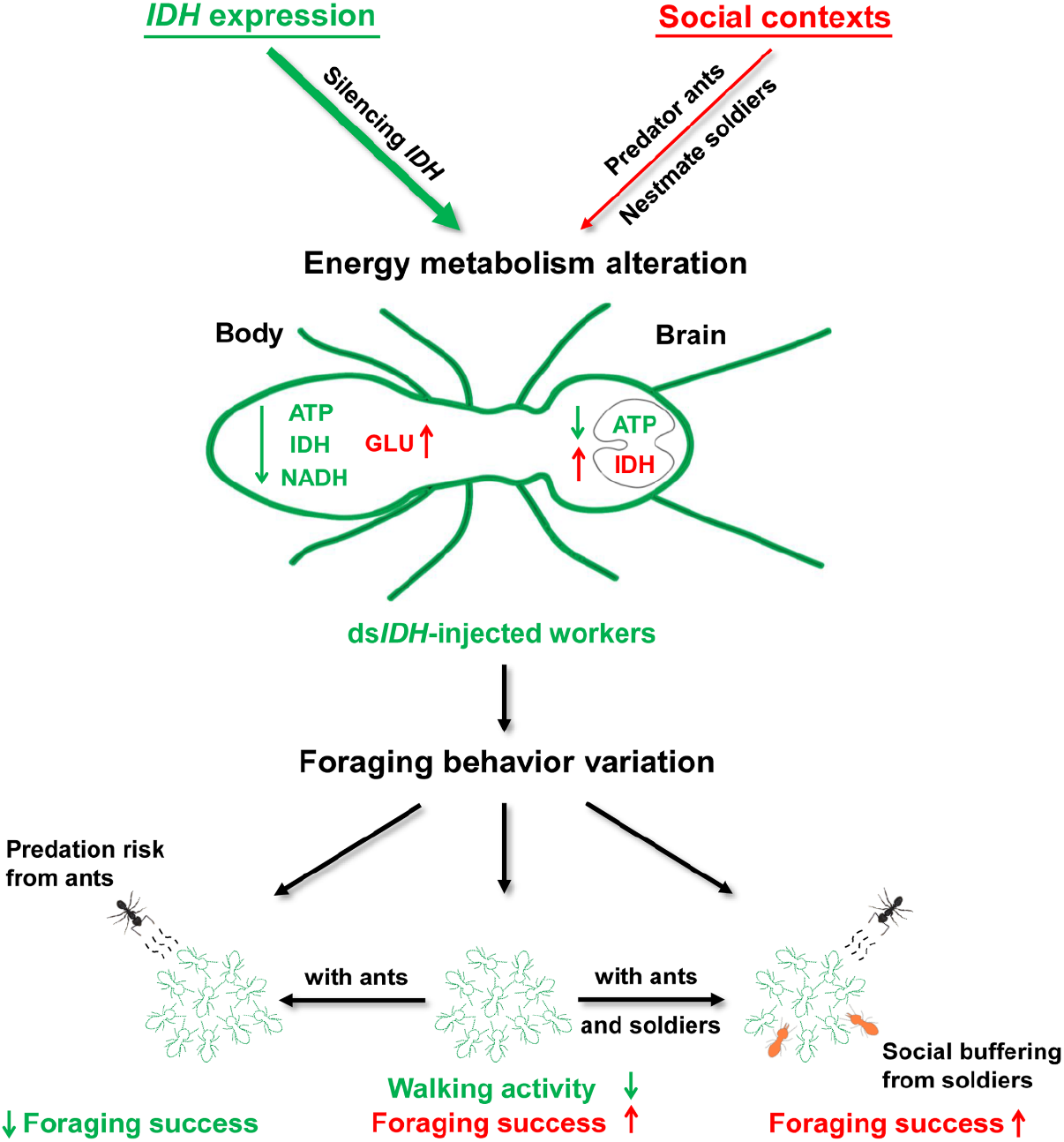
The role of the IDH gene and the social context in regulating the foraging behavior of *O. formosanus*. *IDH* silencing impaired the NAD^+^-IDH reaction in the TCA, leading to a decrease in ATP levels, IDH activity and the NADH levels but an increase in glucose levels in the whole organism, resulting in increased IDH activity but decreased the ATP level in the brain. When ds*IDH*-injected workers foraged together, their velocity and distance moved decreased, but their frequency and cumulative duration in food zones increased, suggesting that *IDH* downregulation reduced walking activity but enhanced foraging success. The social context could also alter the brain energy metabolism of foraging workers, including decreasing ATP levels but the increasing IDH activity in the social context with ants and soldiers, which further changed the foraging behavior of the workers. When predator ants were present, the ds*IDH*-injected workers decreased their frequency and cumulative duration in food zones, showing a significant decline in foraging success. However, the increase in the number of nestmate soldiers strengthened social buffering to relieve the negative effect of predator ants on worker foraging behavior and, thus, improved the foraging success of ds*IDH*-injected workers. Our orthogonal experiments verified that the role of the IDH gene as an inherent factor was dominant in modulating termite foraging behavior compared to the external social context (predator ants and nestmate soldiers). Thus, abnormal energy metabolism mediated by *IDH* altered termite foraging behavior in different social contexts.

## Materials and Methods

### Experimental termites

We collected termite samples from 26 *O. formosanus* colonies in the field (Table S1). Samples of ants were collected as predators from a single *L. kitteli* colony in the field. All the termite and ant colonies were collected from Shizi Hill, Wuhan City, China. They were raised under controlled laboratory conditions (darkness, 25 ± 1 °C, 80 ± 5% relative humidity). Healthy termite and ant samples were chosen for the subsequent experiments.

### Cloning and sequencing of the IDH gene

The termite samples were frozen in liquid nitrogen immediately after collection. RNA was extracted with RNAiso Plus (Takara Bio Inc., Tokyo, Japan), and cDNA was synthesized using a PrimeScript RT reagent kit with gDNA Eraser (Takara Bio Inc.). Gene-specific primers for the complete ORF of *IDH* were designed based on partial sequences of Unigene 32825 obtained from the transcriptome data of the heads of *O. formosanus* workers [48]. The nested polymerase chain reaction (PCR) amplification reactions were carried out as follows: 95 °C for 4 min; 38 cycles of 95 °C for 30 s, 55 °C for 30 s and 72 °C for 80 s; and one cycle at 72 °C for 5 min. Following PCR amplification, the *IDH* fragment was cloned, purified and sequenced.

### Sequence and phylogenetic analyses of the IDH gene

Sequence alignment and analysis were performed using the BLAST service of the National Center for Biotechnology Information (NCBI, http://www.ncbi.nlm.nih.gov). Protein multiple alignment analyses were performed using MEGA 6.0 and GeneDoc 2.0 software. The deduced amino acid sequence of *IDH* was aligned with its corresponding orthologs from the harvester ant *Pogonomyrmex barbatus* (XP_011645897.1), the honey bee *Apis mellifera* (XP_006564183.2) and *Homo sapiens* (NP_005521.1). cDNA and amino acid sequence similarity searches were performed using the BLAST algorithm (https://blast.ncbi.nlm.nih.gov/Blast.cgi). The phylogenetic relationships of *IDH* and the homologous genes in other species were analyzed using the neighbor-joining (NJ) method with MEGA 6.0 software with 1,000 bootstrapping replicates. The prediction of protein secondary structure was performed using the Self-Optimized Prediction Method with Alignment algorithm of the online software PRABI-Lyon-Gerland (https://prabi.ibcp.fr/htm/site/web/home).

### *IDH* transcription in different tissues of *O. formosanus*

The head, abdomen and thorax tissues were dissected from *O. formosanus* workers under low-temperature conditions. Thirty individual thorax specimens and 15 individual head and abdomen specimens were prepared for each replicate. RNA extraction and cDNA synthesis from these body regions were carried out as described in the section on the cloning and sequencing of the IDH gene. Quantitative real-time PCR (qRT-PCR) was performed with Hieff™ qPCR SYBR^®^ Green Master Mix (Yeasen, China) in a QuantStudio 6&7 Flex Real Time PCR System (Applied Biosystems, Life Technologies, Milan, Italy). The relative expression levels of *IDH* among the three body regions were calculated using the 2^−ΔΔCt^ method [71]. Six biological replicates were performed for the RT-qPCR analysis of *IDH*. The primers used for RT-qPCR are listed in Table S2.

### RNA interference with *IDH* transcription

The template cDNAs were amplified by using PCR primers that had T7 RNA polymerase sequences appended to their 5’ ends. The PCR primers for the dsRNA template (shown 5’ to 3’) are shown in Table S2. Injection was performed using a sterilized microinjector (SYS-PV820, World Precision Instruments, USA). Approximately 3 μg of dsRNA targeting *IDH* or *GFP*, or 150 nl of water was injected into the side of the thorax [72]. After injection, the termites were transferred to petri dishes with filter paper containing a 7.5% glucose solution. Individuals and brain tissues were collected separately at 3 d after injection for subsequent experiments. qPCR was performed for each sample using a QuantStudio 6&7 Flex Real Time PCR System. Then, cDNA to be used as the template for qRT-PCR was synthesized from the total RNA of each of six individuals at 3 d after injection. Gene-specific primers were designed by using NCBI primer-BLAST and are presented in Table S2. mRNA levels were quantified using *β-actin* and *glyceraldehyde-3-phosphate dehydrogenase* (*GAPDH*) as the reference genes. The relative expression levels of specific genes were calculated via the 2^−ΔΔCt^ method [71]. Each treatment consisted of at least 6 biological replicates.

### Brain tissue anatomy and immunocytochemistry with synapsin

Termite brains were dissected out in an ice-cold sterilized 0.9% sodium chloride solution in a dissecting dish cooled on ice [73] and then immediately frozen with liquid nitrogen for *IDH* expression and metabolite assays. The insects were anesthetized by cooling on ice. The brains were dissected out and fixed in a 4% paraformaldehyde solution in phosphate-buffered saline (PBS; in mM, 684 NaCl, 13 KCl, 50.7 Na_2_HPO_4_, 5 KH_2_PO_4_, pH 7.4) for 2 h at room temperature. After fixation, the brains were rinsed in PBS for 4 × 15 min. To minimize nonspecific staining, the brains were preincubated in 5% normal goat serum (Sigma, St. Louis, MO) in PBS containing 0.5% Triton X-100 (PBSX; 0.1 M, pH 7.4) for 3 h at room temperature. Then, the brains were incubated in the primary antibody SYNORF1 at 1:100 in PBSX at 4 °C for 3 d. After incubation, the brains were rinsed in PBS for 4 × 15 min before being incubated in the Cy2-conjugated anti-mouse secondary antibody (dilution 1:300 in PBSX; Invitrogen, Eugene, OR), at 4 °C for 1 d. The brains were finally rinsed for 4 × 15 min in PBS, dehydrated in an ascending ethanol series, and mounted in methyl salicylate [74]. A confocal laser scanning microscope (LSM 510, META Zeiss, Jena, Germany) was used to scan the brain and to obtain serial optical images with excitation by a 488 nm laser.

### Metabolite assays

Three days post-injection, termite individuals were weighed and then immediately crushed in liquid nitrogen and dissolved at a ratio of 10 mg of body fresh weight to 100 μL of the solution recommended by the manufacturer. Termite brains were dissected 15 min after foraging observations in different social contexts and then dissolved in the solution recommended by the manufacturer. The determination of the ATP (n = 9 replicates), NADH (n = 6 replicates), IDH (n = 6 replicates) and glucose (n = 9 replicates) contents of the whole organism and the ATP (n = 6 replicates) and IDH (n = 6 replicates) contents of the brain were performed according to the protocols provided by the manufacturers (Beyotime Biotechnology (Shanghai, China), Nanjing Jiancheng Bioengineering Institute (Jiangsu, China) and Zeye Biological Technology (Shanghai, China)). The protein concentration of the brain was determined according to the protocol of the BCA (bicinchoninic acid) protein concentration determination kit from Beijing Dingguo Changsheng Biotechnology (Beijing, China).

### Termite behavioral apparatus

The circular behavioral apparatus included one inner ring (D = 300 mm) in which 10 termites tested and a 10 mm outer ring containing the predator ant *L. kitteli*. Both rings of the testing arena had a 10 mm-deep wall edge (Fig 3A). The wall of the inner ring included 40 evenly spaced 1 mm slits that allowed the transmission of chemical cues and antennal contacts but were too narrow for lethal interactions [24]. Four pieces of 20 mm circular filter paper moistened with 7.5% glucose solution were placed on marked food patches.

### Foraging assays

In the foraging assays performed without regard to the social context (*L. kitteli* predator ants and nestmate soldiers), 10 workers injected with ds*IDH* or ds*GFP* were placed in different 35 mm Petri dishes before the start of the experiment. The foraging assay began after the ten workers were added into the center of the inner ring of the testing arena with moistened filter paper, and the foraging behavior of the 10 workers was then recorded by using the trajectory tracking software EthoVision XT (Noldus Information Technology). After recording for 15 min, the tested workers were removed and then quickly placed in liquid nitrogen. Considering the adaptation period of termites in the behavioral apparatus, the features of the foraging behavior of the ten workers were analyzed after 5 min of recording.

In the foraging assays performed in different social contexts (with or without *L. kitteli* predator ants or/and nestmate soldiers), workers and soldiers that did not receive any injection were individually marked with red color on the abdomen by using PX-21 uni-Paint markers prior to the assay so that each individual could be identified. To reduce the potential for injury, each body part of an individual was only marked once [24]. For the convenience of experimental operation, different injected termites were placed in different 35 mm Petri dishes before the start of the experiment. For example, the 24 ds*IDH*-injected termites were transferred to three 35 mm Petri dishes, and each Petri dish contained eight termites. Similarly, the 24 termites that were injected with ds*GFP* or water were transferred to Petri dishes respectively. Each Petri dish was covered with filter paper moistened with water. According to orthogonal experimental design L_9_ (3^3^) (Table 1), we transferred one or two marked workers or/and marked soldiers that were not subjected to dsRNA or water injection separately into nine Petri dishes (Fig 5A-D). Finally, there were a total of 10 termites in each Petri dish, including unmarked and marked individuals.

At the beginning of the experiments, the inner ring of the testing arena was covered with moistened filter paper. According to the orthogonal experimental design L_9_ (3^3^) (Table 1), we placed different numbers of *L. kitteli* predator ant workers in the outer ring of the testing arena for 30 min before adding termites. The foraging assay began after the ten termites (including workers and/or soldiers) were added to the center of the inner ring of the testing arena with moistened filter paper. The features of the behavioral phenotypes of the 10 termites were recorded by using EthoVision XT software. After recording for 15 min, the tested termites were removed and then quickly placed in liquid nitrogen. The features of the foraging behavior of eight workers were analyzed after 5 min of recording, which was also the procedure for the foraging assay without the social context described above.

Four important phenotypic parameters of foraging behavior were measured, including the velocity and distance moved, and the frequency and cumulative duration in food zones. In this study, the velocity and distance moved were used to describe the walking activity of the worker termites, and the frequency and cumulative duration in food zones were used to describe the foraging success of the worker termites [35, 75].

### Statistical analysis

The foraging behaviors of termites were monitored using a digital camera (BASLER, acA1920-40gc) coupled to video tracking software (EthoVision XT 14.0, Noldus Information Technology). All statistical analyses were conducted using IBM SPSS Statistics 19.0 software. The Shapiro-Wilk test was used to verify whether the data conformed to a normal distribution or not. Most of the data showed a normal distribution, except for those for the RNAi efficiency in the brain and the metabolite assays. The expression pattern of *IDH* and the phenotypic parameters of foraging behavior in the orthogonal experiments were analyzed with Tukey’s HSD test. Gene expression, RNAi efficiency in the whole-organism and foraging assays after silencing *IDH* without regard to the social context were analyzed with paired *t*-tests. The impact degrees of three factors (gene, ant number and soldier number) on worker foraging behavior were determined with a general linear model (GLM). The assessment of RNAi efficiency in the brain and metabolite assays was performed with the Wilcoxon test. The significance level in this study was set at *p* < 0.05.

## Acknowledgments

We would like to thank Dr. Zhao Xincheng from Henan Agriculture University for technical help with the brain tissue anatomy and immunocytochemistry of termites.

## Supporting information

**Table S1.** Distribution of the 26 *O. formosanus* colonies for each experiment*

| **Experiments** | **Replicates** | **Colonies** |
| --- | --- | --- |
| Expression pattern of *IDH* | 6 | 1,2,3 |
| RNAi efficiency for *IDH* | 9 | 4,5,6 |
| Survival analysis for *IDH* | 6 | 5,6 |
| Evaluation of IDH enzyme activity | 6 | 7,8 |
| Evaluation of ATP level | 9 | 7,8,9 |
| Evaluation of NADH level | 6 | 10,11 |
| Evaluation of glucose level | 9 | 10,11,12 |
| Evaluation of IDH enzyme activity in the brain | 6 | 13,14,15 |
| Evaluation of ATP level in the brain | 6 | 13,14,15 |
| RNAi efficiency for *IDH* in the brain | 6 | 16,17 |
| Foraging behavior for *IDH* | 10 | 17,18,19,20 |
| Orthogonal experiments for foraging assays | 12 | 21,22,23,24,25,26 |

\* All the termite samples were collected from 26 colonies of *O. formosanus* on Shizi Hill, Wuhan City, Hubei Province, China.

**Table S2.**
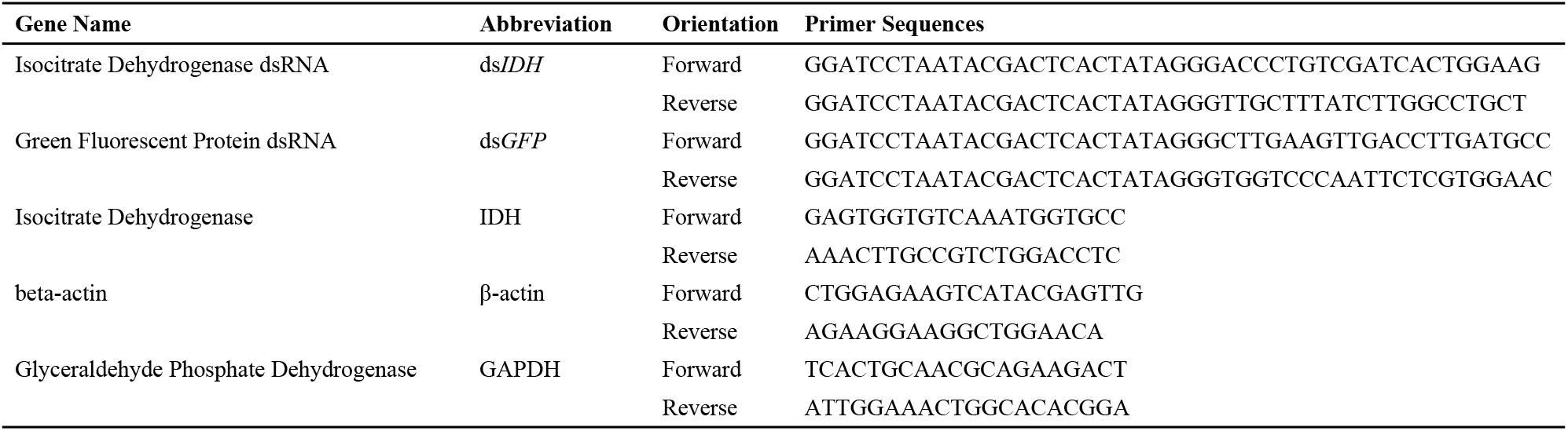
Primers used for qPCR and RNAi

## Reference

1. Mathot KJ, Dingemanse NJ. Energetics and behaviour: unrequited needs and new directions. Trends Ecol Evol. 2015; 30(4): 199–206. http://doi.10.1016/j.tree.2015.01.010 PMID: 25687159

2. Sandhu P, Shura O, Murray RL, Guy C. Worms make risky choices too: the effect of starvation on foraging in the common earthworm (*Lumbricus terrestris*). Can J Zool. 2018; 96(11): 1278–1283. http://doi.10.1139/cjz-2018-0006

3. Camazine S, Deneubourg JL, Franks NR, Sneyd J, Theraulaz G, Bonabeau E. 2001. Self-organization in biological systems. Princeton University Press, Princeton, NJ.

4. Sumpter D, Pratt S. A modelling framework for understanding social insect foraging. Behav Ecol Sociobiol. 2003; 53: 131–144. https://doi.org/10.1007/s00265-002-0549-0

5. Barron AB, Plath J. The evolution of honey bee dance communication: a mechanistic perspective. J Exp Biol. 2017; 220(23): 4339–4346. http://doi.10.1242/jeb.142778 PMID: 29187616

6. Warburg I, Whitford WG, Steinberger Y. Colony size and foraging strategies in desert seed harvester ants. J Arid Environ. 2017; 145: 18–23. http://doi.10.1016/j.jaridenv.2017.04.016

7. Oberst S, Lai JCS, Evans TA. Termites utilise clay to build structural supports and so increase foraging resources. Sci Rep-UK. 2016; 6: 20990. http://doi.10.1038/srep20990 PMID: 26854187

8. Yeates LC, Williams TM, Fink TL. Diving and foraging energetics of the smallest marine mammal, the sea otter (*Enhydra lutris*). J Exp Biol. 2007; 210 (Pt 11): 1960–1970. http://doi.10.1242/jeb.02767 PMID: 17515421

9. Plaçais PY, Preat T. To favor survival under food shortage, the brain disables costly memory. Science. 2013; 339(6118): 440–442. http://doi.10.1126/science.1226018 PMID: 23349289

10. Haagensen AMJ, Sørensen DB, Sandøe P, Matthews LR, Birck MM, Fels JJ, et al. High fat, low carbohydrate diet limit fear and aggression in göttingen minipigs. Plos One. 2014; 9(4): e93821. http://doi.10.1371/journal.pone.0093821 PMID: 24740321

11. Balaskó M, Soós S, Székely M, Pétervári E. Leptin and aging: review and questions with particular emphasis on its role in the central regulation of energy balance. J Chem Neuroanat. 2014; 61: 248–255. https://doi.org/10.1016/j.jchemneu.2014.08.006 PMID: 25218974

12. Feinerman O, Traniello JF. Social complexity, diet, and brain evolution: modeling the effects of colony size, worker size, brain size, and foraging behavior on colony fitness in ants. Behav Ecol Sociobiol. 2016; 70(7): 1063–1074. http://doi.10.1007/s00265-015-2035-5

13. Volkenhoff A, Weiler A, Letzel MC, Stehling M, Klämbt C, Schirmeier S. Glial glycolysis is essential for neuronal survival in *Drosophila*. Cell Metab, 2015; 22(3): 437–447. http://doi.10.1016/j.cmet.2015.07.006 PMID: 26235423

14. Gordon DG, Ilie I, Traniello JFA. Behavior, brain, and morphology in a complex insect society: trait integration and social evolution in the exceptionally polymorphic ant *Pheidole rhea*. Behav Ecol Sociobiol. 2017; 71: 166. https://doi.org/10.1007/s00265-017-2396-z

15. Rittschof CC, Schirmeier S. Insect models of central nervous system energy metabolism and its links to behavior. Glia. 2018; 66(6): 1160–1175. http://doi.10.1002/glia.23235 PMID: 28960551

16. Rittschof CC, Grozinger CM, Robinson GE. The energetic basis of behavior: bridging behavioral ecology and neuroscience. Curr Opin Behav Sci. 2015; 6: 19–27. https://doi.org/10.1016/j.cobeha.2015.07.006

17. Rittschof CC, Vekaria HJ, Palmer JH, Sullivan PG. Brain mitochondrial bioenergetics change with rapid and prolonged shifts in aggression in the honey bee, *Apis mellifera*. J Exp Biol. 2018; 221(Pt 8): jeb176917. http://doi.10.1242/jeb.176917 PMID: 29496782

18. Lima SL. Initiation and termination of daily feeding in dark-eyed juncos: influences of predation risk and energy reserves. Oikos, 1988; 53(1): 3–11. http://doi.10.2307/3565656

19. Nonacs P, Dill LM. Mortality risk vs. food quality trade-offs in a common currency: ant patch preferences. Ecology. 1990; 71(5): 1886–1892. https://doi.org/10.2307/1937596

20. Krams IA. Length of feeding day and body weight of great tits in single- and a two-predator environment. Behav Ecol Sociobiol. 2000; 48: 147–153. https://doi.org/10.1007/s002650000214

21. Fraser DF, Gilliam JF, Akkara JT, Albanese BW, Snider SB. Night feeding by guppies under predator release: Effects on growth and daytime courtship. Ecology. 2004; 85(2): 312–319. https://doi.org/10.1890/03-3023

22. Vijayan S, Kotler BP, Tov-Elem LT, Abramsky Z. Effect of predation risk on microhabitat use by goldfish. Ethol Ecol Evol. 2019; 31(1): 1–11. https://doi.org/10.1080/03949370.2018.1477837

23. Frank ET, Eduard LK. Saving the injured: evolution and mechanisms. Commun Integr Biol. 2017; 10(5-6): e1356516. http://doi.10.1080/19420889.2017.1356516 PMID: 29260800

24. Tian L, Preisser EL, Haynes KF, Zhou XG. Social buffering in a eusocial invertebrate: termite soldiers reduce the lethal impact of competitor cues on workers. Ecology. 2017; 98(4): 952–960. http://doi.10.1002/ecy.1746 PMID: 28122113

25. Hughes NK, Kelley JL, Banks PB. Dangerous liaisons: the predation risks of receiving social signals. Ecol Lett. 2012; 15(11): 1326–1339. http://doi.10.1111/j.1461-0248.2012.01856.x PMID: 22925009

26. Oberst S, Bann G, Lai JC, Evans TA. Cryptic termites avoid predatory ants by eavesdropping on vibrational cues from their footsteps. Ecol Lett. 2017; 20(2): 212–221. http://doi.10.1111/ele.12727 PMID: 28111901

27. Readera T, Higginsona AD, Gilberta BFS. The effects of predation risk from crab spiders on bee foraging behavior. Behav Ecol, 2006; 17(6): 933–939. http://doi.10.1093/beheco/arl027

28. Josens R, Mattiacci A, Lois-Milevicich J, Giacometti A. Food information acquired socially overrides individual food assessment in ants. Behav Ecol Sociobiol. 2016; 70(12): 2127–2138. http://10.1007/s00265-016-2216-x

29. Tian L, Zhou XG. The soldiers in societies: defense, regulation, and evolution. Int J Biol Sci. 2014; 10(3): 296–308. http://doi.10.7150/ijbs.6847 PMID: 24644427

30. Lucas C, Brossette L, Lefloch L, Dupont S, Christidès JP, Bagnères, AG. When predator odour makes groups stronger: effects on behavioural and chemical adaptations in two termite species. Ecol Entomol. 2018; 43(4): 513–524. https://doi.org/10.1111/een.12529

31. Sun PD, Li GH, Jian JB, Liu L, Chen JH, et al. Transcriptomic and functional analyses of phenotypic plasticity in a higher termite, *Macrotermes barneyi* Light. Front Genet. 2019; 10: 964. http://doi.10.3389/fgene.2019.00964 PMID: 31681415

32. Hennessy MB, Kaiser S, Sachser N. Social buffering of the stress response: diversity, mechanisms, and functions. Front Neuroendocrin. 2009; 30(4): 470–482. http://doi.10.1016/j.yfrne.2009.06.001 PMID: 19545584

33. Merchant A, Song DY, Yang XW, Li XR, Zhou XG. Candidate foraging gene orthologs in a lower termite, *Reticulitermes flavipes*. J Exp Zool B Mol Dev Evol. 2020; 334(3): 168–177. http://doi.10.1002/jez.b.22918 PMID: 31713321

34. Sokolowski MB. Foraging strategies of *Drosophila melanogaster*: a chromosomal analysis. Behav Genet, 1980; 10(3): 291–302. http://doi.10.1007/BF01067774 PMID: 6783027

35. Anreiter I, Kramer JM, Sokolowski MB. Epigenetic mechanisms modulate differences in *Drosophila* foraging behavior. P Natl Acad Sci USA. 2017; 114(47): 12518–12523. http://doi.10.1073/pnas.1710770114 PMID: 29078350

36. Ben-Shahar Y, Robichon A, Sokolowski MB, Robinson GE. Influence of gene action across different time scales on behavior. Science, 2002; 296(5568): 741–744. http://doi.10.1126/science.1069911 PMID: 11976457

37. Kodaira Y, Ohtsuki H, Yokoyama J, Kawata M. Size-dependent *foraging* gene expression and behavioral caste differentiation in *Bombus ignitus*. BMC Res Notes, 2009; 2: 184. http://doi.10.1186/1756-0500-2-184 PMID: 19758422

38. Ingram KK, Kleeman L, Peteru S. Differential regulation of the *foraging* gene associated with task behaviors in harvester ants. BMC Ecol. 2011; 11: 19. http://doi.10.1186/1472-6785-11-19 PMID: 21831307

39. Lucas C, Sokolowski MB. Molecular basis for changes in behavioral state in ant social behaviors. Proc Natl Acad Sci USA. 2009; 106(15): 6351–6356. http://doi.10.1073/pnas.0809463106 PMID: 19332792

40. Lucas C, Nicolas M, Keller L. Expression of *foraging* and *Gp-9* are associated with social organization in the fire ant *Solenopsis invicta*. Insect Mol Biol. 2015; 24(1): 93–104. http://doi.10.1111/imb.12137 PMID: 25315753

41. Anreiter I, Vasquez OE, Allen AM, Sokolowski MB. Foraging path-length protocol for *Drosophila melanogaster* larvae. J Vis Exp. 2016; (110): 53980. https://doi.org/10.3791/53980 PMID: 27167330

42. Burns JG, Svetec N, Rowe L, Mery F, Dolan MJ, Boyce WT, Sokolowski MB. Gene-environment interplay in *Drosophila melanogaster*: Chronic food deprivation in early life affects adult exploratory and fitness traits. P Natl Acad Sci USA. 2012; 109 Suppl 2 (Suppl 2): 17239–17244. http://doi.10.1073/pnas.1121265109 PMID: 23045644

43. Kohn NR, Reaume CJ, Moreno C, Burns JC, Sokolowski MB, Mery F. Social environment influences performance in a cognitive task in natural variants of the *foraging* gene. PLoS One. 2013; 8(12): e81272. http://doi.10.1371/journal.pone.0081272 PMID: 24349049

44. Wen P, Ji BZ, Sillam-Dussès D. Trail communication regulated by two trail pheromone components in the fungus-growing termite *Odontotermes formosanus* (Shiraki). Plos one. 2014; 9(3): e90906. http://doi.10.1371/journal.pone.0090906 PMID: 24670407

45. Chiu CI, Yeh HT, Li PL, Kuo CY, Tsai MJ, Li HF. Foraging phenology of the fungus-growing termite *Odontotermes formosanus* (Blattodea: Termitidae). Environ Entomol. 2018; 47(6): 1509–1516. http://doi.10.1093/ee/nvy140 PMID: 30239668

46. Huang QY, Lei CL, Xue D. Field evaluation of a fipronil bait against subterranean termite *Odontotermes formosanus* (Isoptera: Termitidae). J Econ Entomol. 2006; 99(2): 455–461. http://doi.10.1603/0022-0493-99.2.455 PMID: 16686147

47. Cornelius ML, Osbrink WLA. Effect of Soil Type and Moisture Availability on the Foraging Behavior of the Formosan Subterranean Termite (Isoptera: Rhinotermitidae). J Econ Entomol. 2010; 103(3): 799–807. http://doi.10.1603/ec09250 PMID: 20568626

48. Huang QY, Sun PD, Zhou XG, Lei CL. Characterization of head transcriptome and analysis of gene expression involved in caste differentiation and aggression in *Odontotermes formosanus* (Shiraki). Plos One. 2012; 7(11): e50383. http://doi.10.1371/journal.pone.0050383 PMID: 23209730

49. Xu H, Zhu QK, Hu W, Kim K, Lei CL, et al. Impacts of different environmental factors on tunnelling behaviour of the subterranean termite *Odontotermes formosanus* (Shiraki). Ethol Ecol Evol. 2019; 31(3): 231–239. http://doi.10.1080/03949370.2018.1561524

50. Briffa M, Elwood RW. Use of energy reserves in fighting hermit crabs. Proc Biol Sci. 2004; 271(1537): 373–379. http://doi.10.1098/rspb.2003.2633 PMID: 15101696

51. Nation JL. 2015. Insect physiology and biochemistry, 3rd edn. CRC Press, Boca Raton.

52. Liu L, Wang CC, Zhao XY, Guan JX, Lei CL, Huang· QY. Isocitrate dehydrogenase-mediated metabolic disorders disrupt active immunization against fungal pathogens in eusocial termites. J Pest Sci. 2020; http://doi.10.1007/s10340-019-01164-y

53. Matsumura K, Miyatake T. Costs of walking: differences in egg size and starvation resistance of females between strains of the red flour beetle (*Tribolium castaneum*) artificially selected for walking ability. J Evolution Biol. 2018; 31(11): 1632–1637. http://doi.10.1111/jeb.13356 PMID: 30055064

54. Zhang D, Wang Y, Shi Z, Liu J, Sun P, Hou X et al. Metabolic reprogramming of cancer-associated fibroblasts by *IDH3α* downregulation. Cell Rep. 2015; 10(8): 1335–1348. http://doi.10.1016/j.celrep.2015.02.006 PMID: 25732824

55. Mergenthaler P, Lindauer U, Dienel GA, Meisel A. Sugar for the brain: the role of glucose in physiological and pathological brain function. Trends Neurosci. 2013; 36(10): 587–597. http://doi.10.1016/j.tins.2013.07.001 PMID: 23968694

56. Lunt SY, Vander Heiden MG. Aerobic glycolysis: meeting the metabolic requirements of cell proliferation. Annu Rev Cell Dev Biol. 2011; 27: 441–464. http://doi.10.1146/annurev-cellbio-092910-154237 PMID: 21985671

57. Chandrasekaran S, Rittschof CC, Djukovic D, Gu H, Raftery D et al. Aggression is associated with aerobic glycolysis in the honey bee brain. Genes Brain Behav. 2015; 14(2): 158–166. http://doi.10.1111/gbb.12201 PMID: 25640316

58. Niven JE, Laughlin SB. Energy limitation as a selective pressure on the evolution of sensory systems. J Exp Biol. 2008; 211: 1792–1804. http://doi.10.1242/jeb.017574 PMID: 18490395

59. Robinson GE, Fernald RD, Clayton DF. Genes and social behavior. Science. 2008; 322(5903): 896–900. http://doi.10.1126/science.1159277 PMID: 18988841

60. Couzin ID. Collective cognition in animal groups. Trends Cogn Sci. 2009; 13(1): 36–43. http://doi.10.1016/j.tics.2008.10.002 PMID: 19058992

61. Marshall JAR, Bogacz R, Dornhaus A, Planqué R, Kovacs T, Franks NR. On optimal decision-making in brains and social insect colonies. J R Soc Interface. 2009; 6(40): 1065–1074. http://doi.10.1098/rsif.2008.0511 PMID: 19324679

62. White SA, Nguyen T, Fernald RD. Social regulation of gonadotropin-releasing hormone. J Exp Biol. 2002; 205: 2567–2581. PMID: 12151363

63. Goodson JL, Kabelik D, Schrock SE. Dynamic neuromodulation of aggression by vasotocin: influence of social context and social phenotype in territorial songbirds. Biol Lett. 2009; 5(4): 554–556. http://doi.10.1098/rsbl.2009.0316 PMID: 19493876

64. Kamhi JF, Traniello JFA. Biogenic amines and collective organization in a superorganism: neuromodulation of social behavior in ants. Brain Behav Evol. 2013; 82: 220–236. http://doi.10.1159/000356091 PMID: 24281765

65. Li-Byarlay H, Rittschof CC, Massey JH, Pittendrigh BR, Robinson GE. Socially responsive effects of brain oxidative metabolism on aggression. P Natl A Sci. 2014; 111(34): 12533–12537. http://doi.10.1073/pnas.1412306111 PMID: 25092297

66. Hughes JJ, Ward D. Predation risk and distance to cover affect foraging behaviour in Namib Desert gerbils. Anim Behav. 1993; 46(6): 1243–1245. http://doi.10.1006/anbe.1993.1320

67. Thomson JS, Watts PC, Pottinger TG, Sneddon LU. Plasticity of boldness in rainbow trout, *Oncorhynchus mykiss*: do hunger and predation influence risk-taking behaviour? Horm Behav. 2012; 61(5): 750–757. http://doi.10.1016/j.yhbeh.2012.03.014 PMID: 22498695

68. Tamburello N, Côté IM, Dulvy NK. 2015. Energy and the scaling of animal space use. Am Nat. 2015; 186(2): 196–211. http://doi.10.1086/682070 PMID: 26655149

69. Campos-Candela A, Palmer M, Balle S, Alvarez A, Alos J. A mechanistic theory of personality-dependent movement behaviour based on dynamic energy budgets. Ecol Lett. 2019; 22(2): 213–232. http://doi.10.1111/ele.13187 PMID: 30467933

70. Ishikawa Y, Miura T. Hidden aggression in termite workers: plastic defensive behaviour dependent upon social context. Anim Behav. 2012; 83(3): 737–745. https://doi.org/10.1016/j.anbehav.2011.12.022

71. Livak KJ, Schmittgen TD. Analysis of relative gene expression data using realtime quantitative PCR and the 2(-Delta Delta C(T)) Method. 2001; 25(4): 402–408. http://doi.10.1006/meth.2001.1262 PMID: 11846609

72. Zhou X, Oi FM, Scharf ME. Social exploitation of hexamerin: RNAi reveals a major caste-regulatory factor in termites. Proc Natl Acad Sci USA. 2006; 103(12): 4499–4504. http://doi.10.1073/pnas.0508866103 PMID: 16537425

73. Ishikawa Y, Aonuma H, Sasaki K, Miura T. Tyraminergic and octopaminergic modulation of defensive behavior in termite soldier. Plos One. 2016; 11(5): e0154230. http://doi.10.1371/journal.pone.0154230. PMID: 27196303

74. Zhao XC, Ma BW, Berg BG, Xie GY, Tang QB, Guo XR. A global-wide search for sexual dimorphism of glomeruli in the antennal lobe of female and male *Helicoverpa armigera*. Sci Rep-UK. 2016; 6: 35204. http://doi.10.1038/srep35204 PMID: 27725758

75. Hughson BN, Anreiter I, Chornenki NLJ, Murphy KR, Ja WW et al. The adult foraging assay (AFA) detects strain and food-deprivation effects in feeding-related traits of Drosophila melanogaster. J Insect Physiol. 2018; 106(Pt 1): 20–29. http://doi.10.1016/j.jinsphys.2017.08.011 PMID: 28860037

